# When Brands Take a Stand: Evidence from Eye-Movements and Memory Recognition

**DOI:** 10.64898/2026.07.29.741481

**Authors:** Rebecca Oliveri, Yanina Tena Garcia, Katja Fiehler

## Abstract

Brands increasingly express their positions on social issues through advertising, social media, and public campaigns. How such messages shape consumer perception is an ongoing area of investigation. Addressing a gap in brand activism research in marketing, this study uses eye-tracking technology to examine its impact on brand attitude, eye movement behavior, and memory retention. We tested these variables across three experimental tasks: an online survey, a free-viewing task, and a surprise memory recognition test. As a result, disagreement increased reading time for text statements, while agreement increased visual engagement (fixations) with the depicted product. Positive brand attitude also enhanced memory recognition for advertisements. Overall, disagreement appears to trigger effortful processing of conflicting information, while agreement facilitates visual absorption and strengthens product recognition. These findings highlight the cognitive risks and rewards of brand activism and provide a foundation for future research on its underlying perceptual and cognitive mechanisms.

## Introduction

In our everyday lives, we are exposed to thousands of marketing stimuli, ranging from sponsored products and services on social media to eye-catching advertisements in public spaces. In this saturated environment, capturing attention has become a crucial first step in driving brand awareness, purchase decisions, and long-term loyalty. To better connect with consumers, brands are increasingly adopting innovative strategies, among which brand activism has emerged as a prominent approach. Defined as companies taking public stances on social, environmental, or political issues to engage with consumers by aligning with or challenging their values and personal beliefs (1) brand activism has gained popularity over the last decade (2).

While potentially powerful, brand activism comes with risks. Research by Mukherjee and Althuizen (2020) has shown that, while positive consumer-brand agreement does not normally elicit strong reactions, consumer-brand disagreement often triggers negative consumer attitudes. Such asymmetry, evident in the fact that consumer-brand disagreement triggers much stronger negative reactions than the moderate positive responses generated by consumer-brand agreement, poses real reputational risks for companies, as disagreement can erode trust and damage long-term relationships. However, research on brand activism has been limited to survey-based approaches, highlighting a gap that motivated the present study. We thus examine brand attitude, eye movements, and memory recognition to gain a deeper understanding of brand activism’s effects and provide initial insights into the underlying cognitive processes.

## The role of eye movements in brand perception

In order to delve into the cognitive mechanisms underlying consumer reactions to brand activism, it is important to highlight two interrelated processes that shape visual perception: bottom-up (stimulus-driven) and top-down (goal-driven) pathways (3). While the bottom-up pathway is driven by stimulus properties that inherently attract observers’ fixations, such as contrast, color, or motion, independent of the observer’s internal state (4), the top-down pathway reflects cognitive influences that guide eye movements based on the observer’s goals and intentions (4). These two mechanisms interact dynamically through underlying neural systems (5), and the comprehension of this dual-system framework is critical for understanding how visual perception and memory coordinate to optimize perception, decision-making, and action in complex environments. Eye- tracking provides a valuable method for capturing these processes in real time, offering precise measures of how individuals allocate visual attention and process information in brand communications (6–8). Among others, eye-tracking technology offers key metrics such as fixations (6, 9), which represent periods of relative stability of the eyes during which information can be processed (6). Emotional content, in the case of brand activism tied to moral values, can further bias visual perception and processing speed. Previous research has shown that, in the case of emotional images, participants fixated longer during the first 500 ms of stimulus presentation, showing a bias in perceptual focus (10), and an increased number of fixations in response to emotionally charged scenes (11). Usée et al. (2020) further investigated the role of emotionally valid stimuli in an eye-tracking study, assessing emotional impact on reading behavior. Their findings indicated that emotionally positive text stimuli were perceived as more pleasant and less arousing than emotionally negative text stimuli. This difference in perception was reflected in shorter reading times for positively valenced texts compared to negatively valenced texts. These dynamics suggest that consumer agreement or disagreement with a brand’s stance may systematically alter visual behavior.

## The role of memory in brand perception

Alongside capturing attention, brand recognition and recall are critical factors in influencing consumers’ attitudes (12, 13). Visual perception plays a crucial role in memory processes, as it determines which elements of the environment are encoded into memory and how that information is later retrieved and used in decision-making (8, 14). When referring to the recognition of visual stimuli, Visual Long-Term Memory (VLTM) (15) depends on the interplay of top-down and bottom-up processing, although it is more strongly shaped by intentions and expectations through top-down processes (16). Given prior findings that consumer–brand agreement shapes brand attitudes (17), it is essential to understand how these attitudes, in turn, can influence recognition memory for brand activism messages.

## Research aims and hypotheses

Against this background, this study aims to lay the groundwork for assessing how responses to brand activism are reflected in perceptual and cognitive processes. We examined the characteristics of viewers’ eye movements and memory performance as they looked at a sequence of custom-made advertisements created with AI tools. To investigate this, we used a multi-method design that combines surveys, eye-tracking, and memory tasks. The study was guided by three main hypotheses:

H1. Brand attitude is affected by consumer-brand agreement.

H2/A. Consumer-brand agreement influences the time spent looking at the advertisement and brand statement.

H2/B. Consumer-brand agreement influences the fixation frequency of the advertisement and brand statement.

H3. Brand attitude influences memory recognition of advertisements used in brand activism.

By integrating eye-tracking and memory measures with traditional survey data, this study offers new insights into the cognitive risks and rewards of brand activism, advancing theory on consumer information processing while providing real-life implications for brand communication strategies.

## Method

### Participants

Forty-five students from Justus-Liebig-University Giessen (33 female, age range = 18-35 years, M = 22.96, SD = 3.86) took part in the study in exchange for course credit. All reported normal or corrected-to-normal vision and were fluent in English, as the entire set of stimuli and instructions was presented in English. All participants provided written informed consent, and the experiment was conducted in compliance with the guidelines of the local ethical committee of the Justus-Liebig-University Giessen and the Declaration of Helsinki (2013). Since the experiment involved custom-made stimuli, all participants signed a disclaimer (acknowledging the fictional and AI-generated nature of the materials) at the end of the experiment. Data collection started on November 6th 2024 and ended on February 12th 2025.

## Apparatus

Participants were seated on a height-adjustable chair in front of a 25" monitor (1920 x 1080 resolution, 60 Hz refresh rate), with a chin rest to stabilize head position and maintain a consistent distance of 90 cm to the monitor. Eye movements were recorded using the video-based Desktop Mount EyeLink 1000 (SR Research Ltd., Mississauga, Ontario, Canada; sampling rate: 1000 Hz), positioned on an elevated surface without obstructing any part of the monitor. Eye movement accuracy was calibrated and validated with a 5-point grid calibration. Viewing was binocular, though only the right eye movements were recorded.

## Materials

The stimuli consisted of 40 advertisement images and 20 brand statements (1000x1000 pixels) depicting fictitious brands and their products. Each image was structured in the same way and comprised a brand name, a slogan, and a product (Figure 1). The 40 advertisement images were divided into two sets: 20 main images (Figure 1A) were used for the viewing task, where each image was paired with a brand statement; the remaining 20 served as distractor images (Figure 1B) in the memory task (see Supplementary File S3). The image set ranged across diverse product categories: five cosmetics (two perfumes and three skincare products), two home accessories, two footwear items, four food and beverage products (one fast-food brand, two supermarkets, and one beverage), and seven accessories (including two backpacks, one pair of socks, one jewelry item, one watch, and two general accessories).

**Figure 1.**
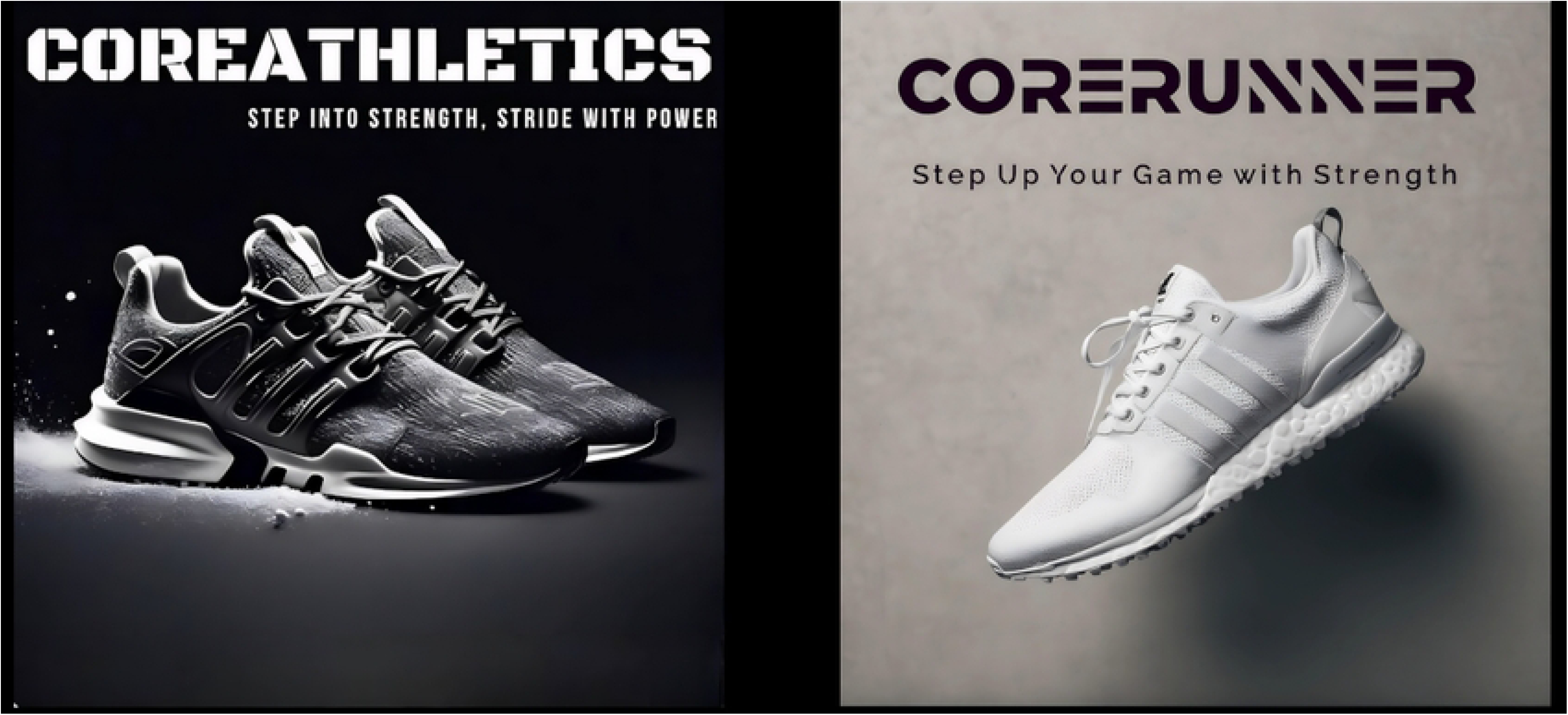
Example Stimuli used in the Viewing and Memory Task. A) Main stimulus presented during the viewing task. B) Paired distractor image presented during the memory task.

The 20 brand statements were created using the language model OpenAI ChatGPT (2023) (see Supplementary File S2) and were designed to mirror an Instagram post template so as to enhance realism and effectively engage participants. Each statement comprised approximately 110 words, including two hashtags, and referenced the name of the fictitious brand. The statements clearly expressed either support for or opposition toward the selected activism topics, including climate change, abortion rights, immigration rights, animal protection, environmental sustainability, gender parity, and LGBTQ+ rights. The statements were balanced, with 50% advocating for a respective topic and 50% opposing it. To assess the perceived positivity or negativity of each statement, a survey (see Supplementary File S4) was distributed to ten additional participants (five female, five male, age range = 24-40 years). Based on survey data, all statements were subsequently refined to ensure clarity and impact.

## Procedure

The experiment consisted of three parts: an online survey, a viewing task, and a surprise memory task. Participants first completed an online survey with 15 questions, starting with demographic information and followed by a question assessing English proficiency (see Supplementary File S1). The remaining questions were all 7-point Likert scale items addressing participants’ positions on social issues (climate change, abortion rights, immigration rights, animal protection, environmental sustainability, gender parity). These topics were selected based on their heightened activity and impact in recent years (2).

Upon completion of the online survey, participants were invited to the university to participate in the viewing and memory tasks of the experiment. Before beginning the experiment, participants were briefed on the viewing task but were not informed about the subsequent memory task. In the viewing task, each trial began with a fixation cross displayed at the center of the screen (see Figure 2). The trial proceeded when participants maintained a stable fixation on this cross for one second. Depending on the presentation order condition the participant was assigned to (*Ad first* vs. *Statement first*), the fixation cross was either followed by an advertisement image or a brand statement. Participants were instructed to view the stimulus at their own pace, without any time constraints, and were allowed to move their eyes freely across the picture. They would then proceed to the next trial by clicking the left button of the computer mouse. This action triggered the reappearance of the fixation cross, which again had to be fixated for one second. Following this, the second stimulus appeared, complementing the first (i.e., brand statement if the ad image was shown first, or vice versa). As before, participants viewed the stimulus without time constraints and advanced by clicking the mouse when ready. After both stimuli were presented, participants were required to rate the brand on three 7-point Likert scales: pleasantness (1 = unpleasant, 7 = pleasant), likeability (1 = dislike, 7 = like), and morality (1 = bad, 7 = good). Ratings and response times were recorded. A total of 20 trials were performed. Brand order presentation was randomized across participants.

**Figure 2.**
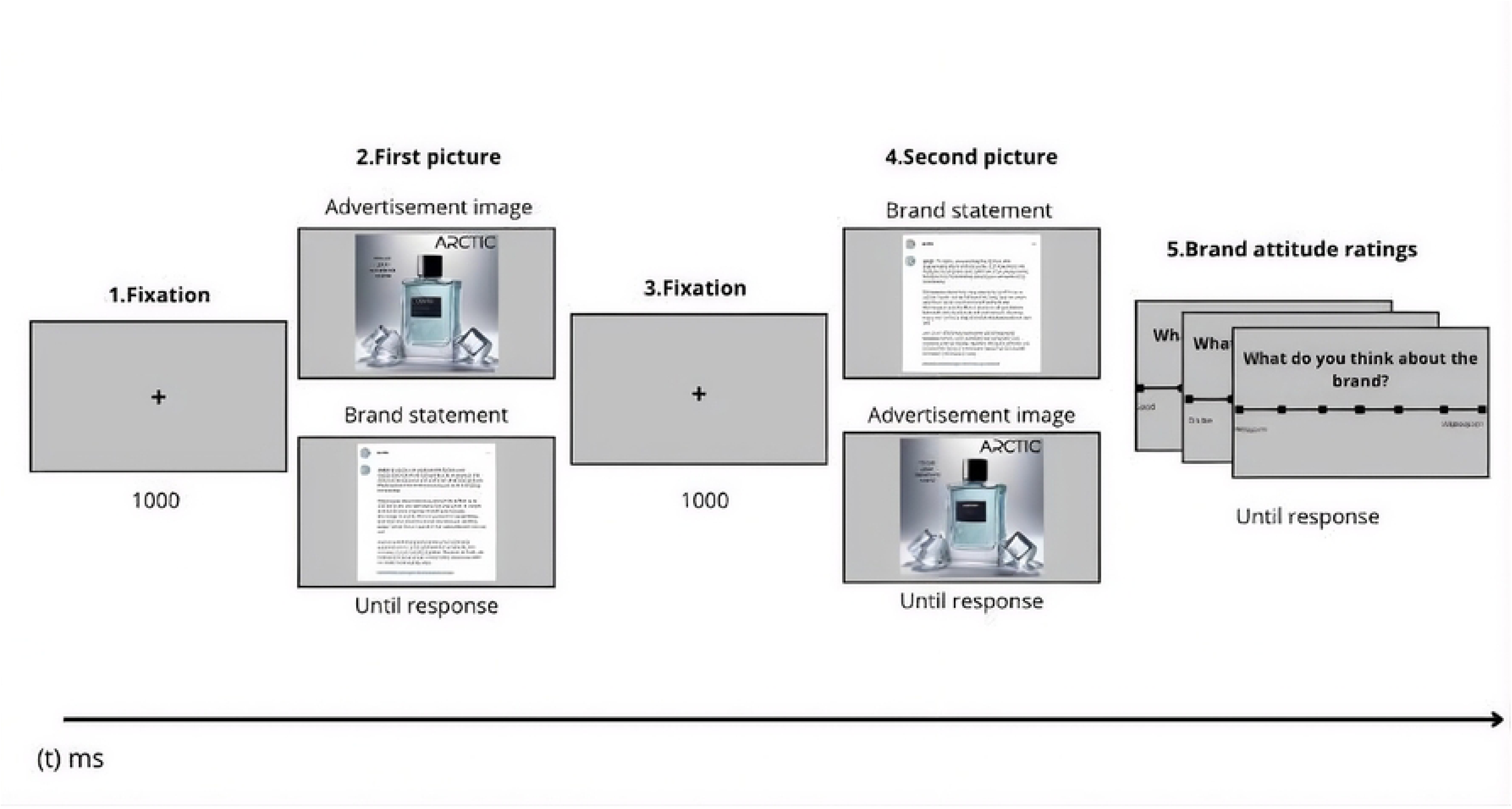
Experimental Procedure.

Following a 5-minute break, participants completed a surprise recognition memory task. In this part, participants were tasked with identifying whether they had seen a given advertisement image before or not (yes/no response). They were informed that reaction times were recorded, so they had to respond as quickly as possible by pressing the "Y" key on the keyboard for indicating "Yes, I have seen this picture before" with their left index finger, and the "N" key for "No, I haven’t seen this picture before" with their right index finger. Participants were shown 40 advertisement images (20 previously seen, 20 distractors) in randomized order. Upon the participant’s response by either pressing the Y or N key, the next image appeared immediately. In addition to reaction time, response accuracy was also recorded in this part of the experiment.

## Eye Movement Measurements

Eye movement behavior was assessed using two measurements: 1) time spent viewing the advertisement and statement image and 2) percentage of fixations in a trial that landed in the predefined Regions of interest in the advertisement pictures. Regions of interest (ROIs) were predefined by manually outlining the product image and the brand name in each stimulus (for an example see Figure 3). A fixation was classified as landing within a ROI when its spatial coordinates fell inside the ROI outline. Fixations that only partially fell within a ROI were nevertheless included.

**Figure 3.**
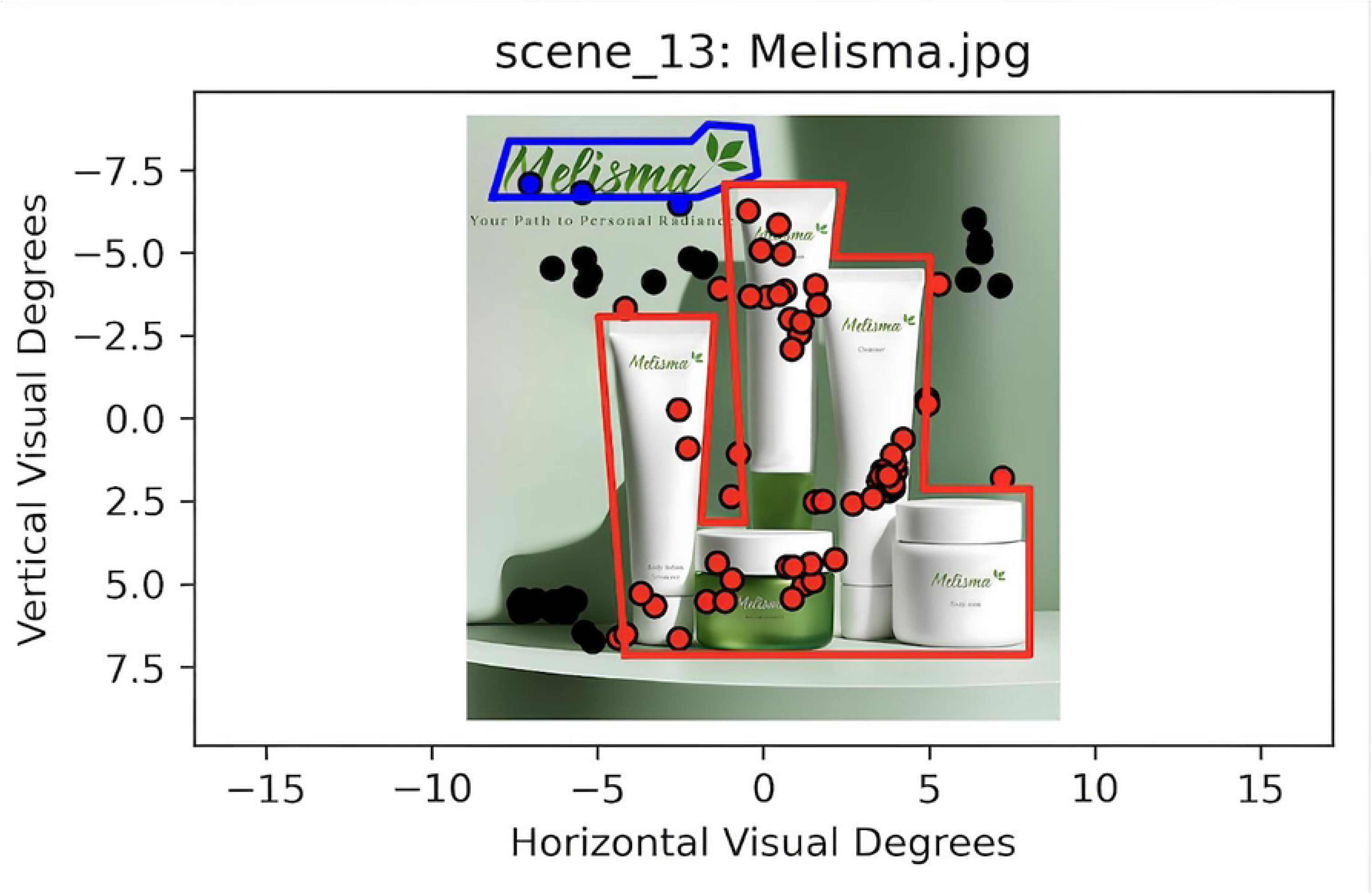
Predefined ROIs. Predefined ROIs, with the product area highlighted in red and the brand name in blue. Participants’ fixations are indicated by dots. Their color indicates whether they were identified as being in a ROI (red meaning they landed on the product and blue on the brand name) or not (black dots).

Eye-tracking data were preprocessed to exclude trials with no registered fixations (10.32% of trials) and participants with > 50% missing trials (n = 5). Blinks longer than 200 ms (20.25% of fixations) or fixations shorter than 15 ms (0.11 %) were also excluded, leaving 34 datasets for statistical analysis.

## Statistical Analyses

All statistical analyses were performed using Jamovi (version 2.6.26). An alpha level of .05 was applied to all analyses, and effect sizes are reported as Cohen’s *d*.

Participants’ responses to the online survey were used to categorize their stance on each activism cause as *pro (*score > 5), *con* (score < 3), or *neutral* (score = 4), based on a 7-point Likert scale. These stances were then compared to the corresponding brand position to determine consumer– brand agreement (agreement, disagreement, neutral). To examine the effect of consumer-brand agreement on brand attitude, a dependent *t*-test was conducted. Only participants showing agreement or disagreement were included in the main analysis and in the memory recognition analysis, resulting in 39 included data sets. Memory performance was assessed with a recognition test and analyzed first for both stimulus types (*original* vs. *new stimuli*) and then for original stimuli only. Dependent *t*-tests examined the effect of stimulus type (*original* vs. *new stimuli*) on reaction times and response accuracy, while brand attitude effects on memory for original stimuli were tested with dependent *t*-tests.

Since consumer–brand agreement could only be formed after the statement was presented, agreement effects were analyzed in the *Statement first* condition only. Accordingly, two separate analyses were performed for 1. the effect of consumer–brand agreement on brand attitude, 2. the effect of consumer–brand agreement on eye movements, and 3. the effect of brand attitude on memory recognition. The first analysis used an independent *t*-tests to assess presentation order effects across the full sample, while the second used a dependent *t*-tests to test the effect of consumer–brand agreement within the *Statement first* condition.

## Results

Data were analyzed in relation to the three research questions: 1. the effect of consumer–brand agreement on brand attitude, 2. the effect of consumer–brand agreement on eye movements, and 3. the effect of brand attitude on memory recognition.

## Brand Attitude

The results revealed a significant difference in consumer-brand agreement on brand attitude [t (38) = 10.6, p < .001, d = 1.70]. As illustrated in Figure 4, mean brand attitude was significantly higher under agreement (M = 5.65, SE = 0.12) compared to disagreement (M = 3.18, SE = 0.14). Relative to a neutral midpoint of 4, these results show a positive brand attitude (score > 4) for consumer- brand agreement and a negative attitude (score < 4) for consumer-brand disagreement, confirming H1.

**Figure 4.**
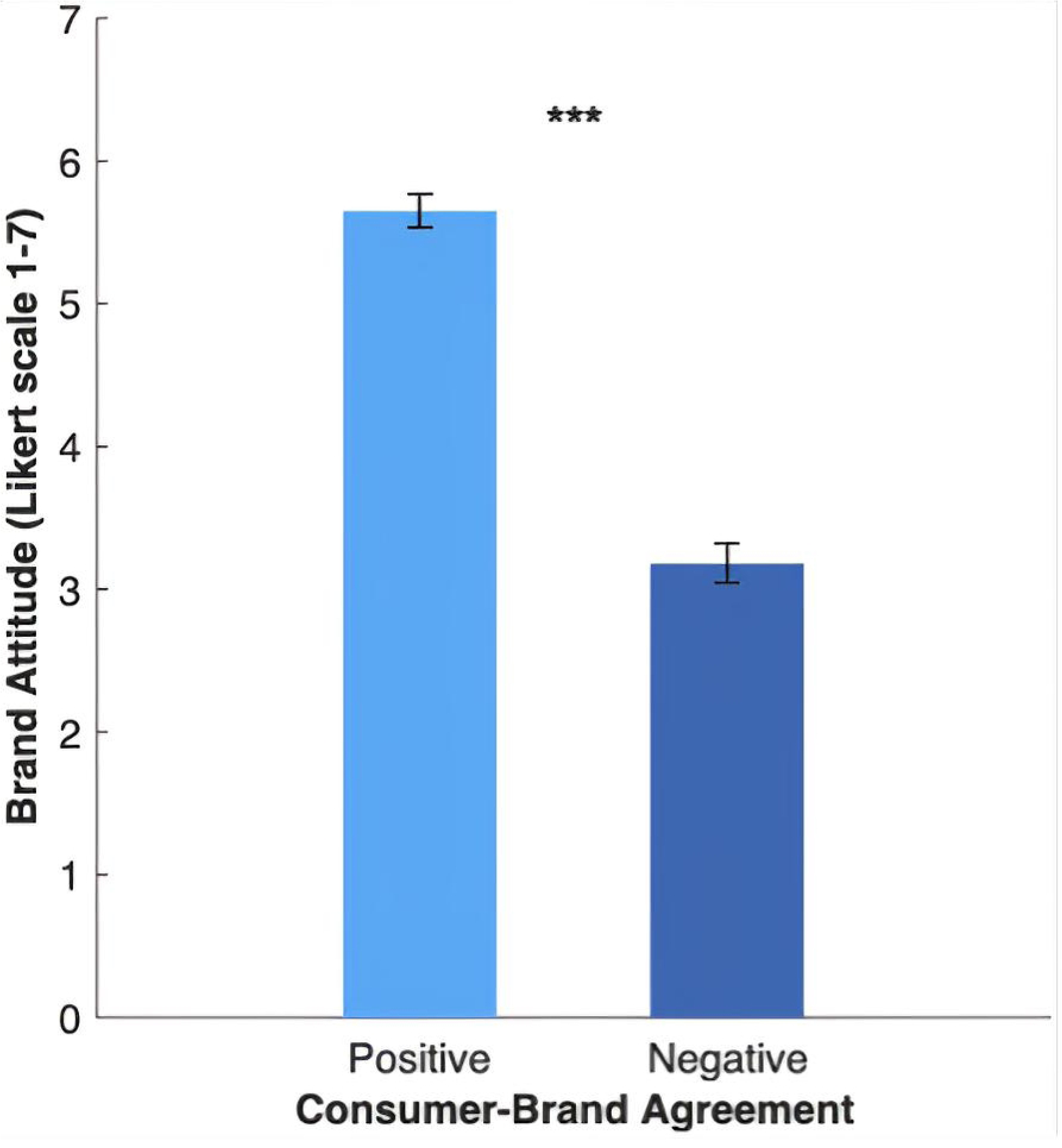
Effect of Consumer-Brand Agreement on Brand Attitude.

## Time Spent on the Advertisement

The results showed a significant difference between the presentation order *Ad first* and *Statement first* [t (32) = 2.19, p = .036, d = 0.752]. As shown in Figure 5A, participants spent significantly more time viewing the image when the advertisement was first (M = 9.62 s, SE = 0.91), compared to when the statement was first (M = 6.78 s, SE = 0.92). However, when examining the main effect of consumer-brand agreement on the time spent viewing the advertisement in the *Statement first* condition, consumer–brand agreement had no effect on viewing time [t (16) = -0.653, p = .523, d = -0.158].

**Figure 5.**
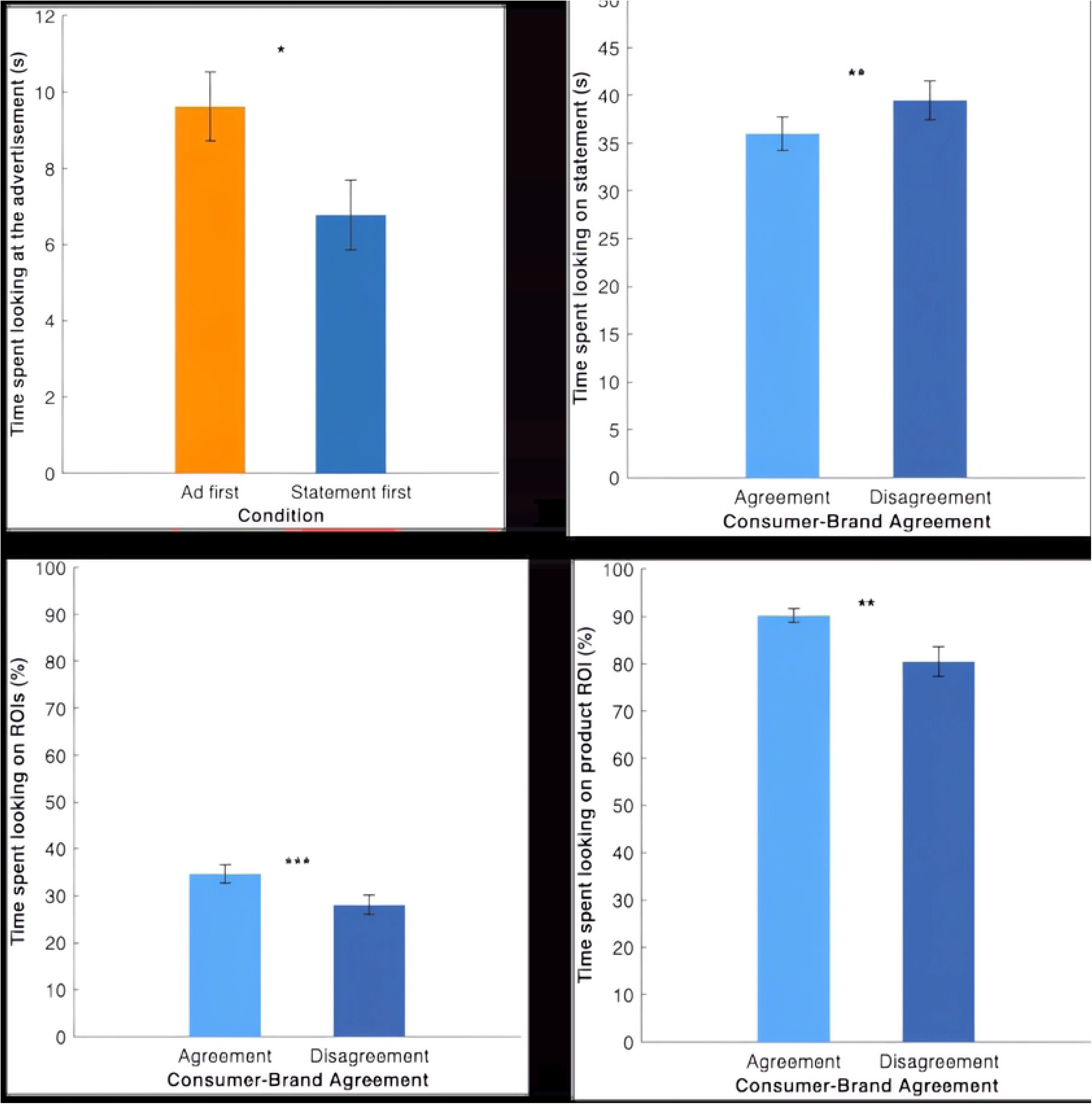
Eye Movement results. A) Effect of Picture Order on Time spent looking on the Advertisement; B) Effect of Consumer-Brand Agreement on Time spent looking on the Statement; C) Effect of Consumer- Brand Agreement on Time spent looking on the ROIs; D) Effect of Consumer-Brand Agreement on Time spent looking on product ROI.

## Time Spent on the Brand Statement

No significant effect of presentation order was found on the time spent viewing the statement [t (32) = 1.81, p = .079, d = 0.622], showing that in both conditions the time needed to read the statement was, on average, about 38 seconds.

However, the analysis of the effect of consumer-brand agreement on time spent viewing the statement showed a significant difference [t (33) = -3.03, p = .005, d = -0.519]. Figure 5B shows that participants spent more time on the statement in case of consumer-brand disagreement (M = 39.5 s, SE = 2.05) compared to consumer-brand agreement (M = 36.0 s, SE = 1.78).

## Fixations on ROIs

Overall, participants allocated ∼32% of trial fixations to ROIs, irrespective of presentation order [t (32) = 1.28, p = .208, d = 0.440]. However, as illustrated in Figure 5C, the analysis conducted to assess the effect of consumer-brand agreement on fixations within the predefined ROIs revealed a significant difference in consumer-brand agreement [t (16) = 4.60, p < .001, d = 1.12]. Participants spent more time on the ROIs in case of consumer-brand agreement (M = 34.7%, SE = 2.02), compared to consumer-brand disagreement (M = 28.1%, SE = 2.09).

Within ROIs, most fixations landed on the product rather than on the brand name (∼85%), with no significant difference between the *Statement first* and *Ad firs*t conditions [t (32) = 0.527, p = .602, d = 0.181]. However, the effect of consumer-brand agreement on time spent on the product ROI revealed a significant difference in consumer-brand agreement [t (16) = 2.94, p = .010, d = 0.714]. Figure 5D indicates that participants fixated more on the product ROIs in case of consumer-brand agreement (M = 90.2%, SE = 1.53) compared to consumer-brand disagreement (M = 80.4%, SE = 3.12).

## Summary

To summarize, presentation order significantly influenced the viewing time on the advertisement, whereas consumer–brand agreement significantly affected viewing time on the statement and directed visual focus toward brand-relevant regions of interest in the advertisement, particularly within the product. These findings support H2/A and H2/B, showing that consumer–brand agreement systematically shapes eye movement behavior.

## Surprise Memory Recognition

### Response Accuracy

As hypothesized, a significant effect was found for stimulus type (Figure 6A), comparing original stimuli with new stimuli [t (38) = 5.09, p < .001, d = 0.815]. Recognition accuracy was higher for original stimuli (M = 94.5%, SE = 0.78) compared to the new distractor images that were seen for the first time (M = 86.1%, SE = 1.59). This lower value for distractors implies a 13.9% false-alarm rate, indicating participants more often mistook new images for ones they had seen before, whereas only 5.5% of the original stimuli were classified as unseen. Further, brand attitude also influenced memory recognition for original stimuli [t (38) = 2.34, p = .024, d = 0.375]. As shown in Figure 6B, participants recognized more advertisements in the case of a positive brand attitude (M = 96.6%, SE = 0.98) compared to a negative brand attitude (M = 92.7%, SE = 1.02). These findings support H3 suggesting that brand attitude enhances recognition memory accuracy.

**Figure 6.**
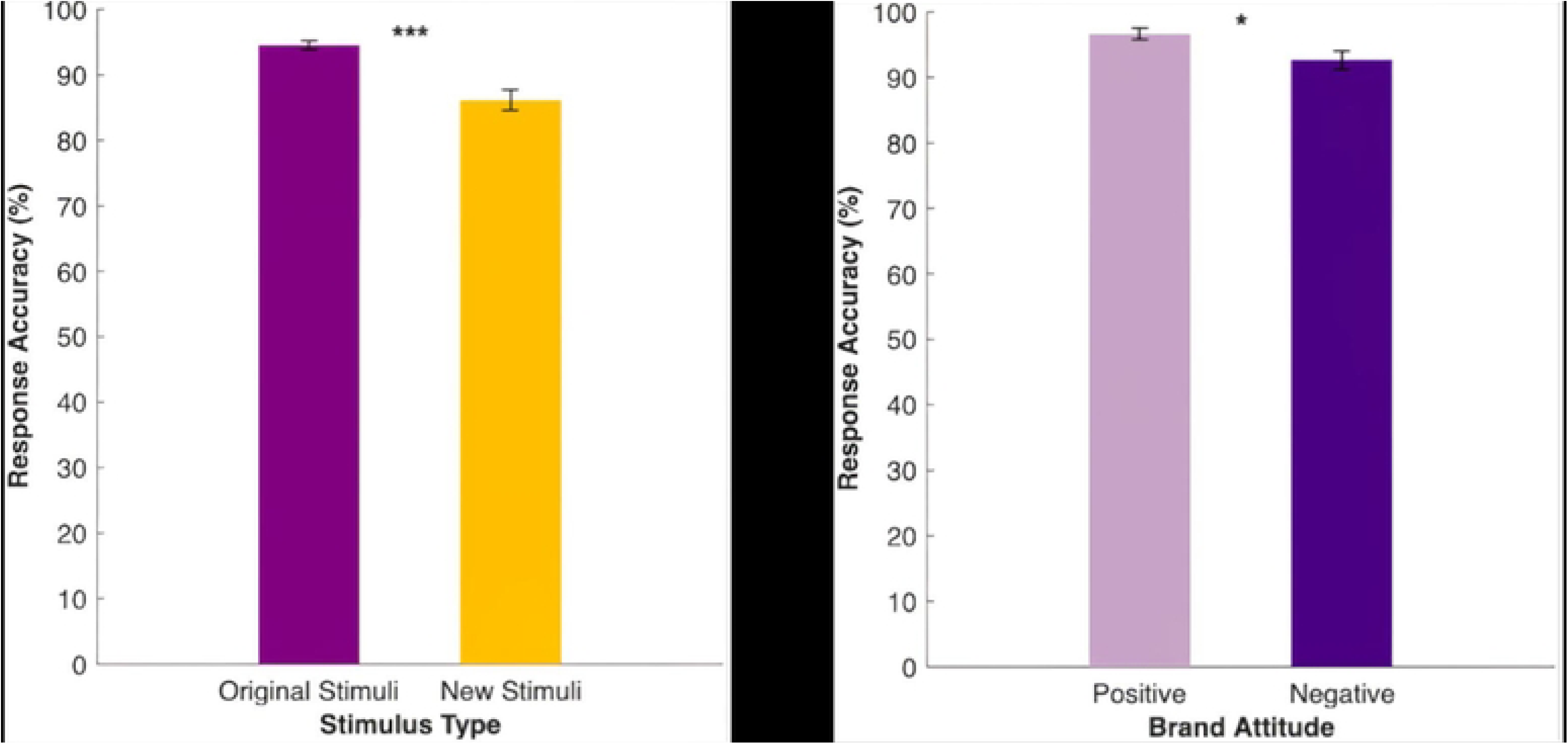
Response Accuracy results. A) Effect of Stimulus Type on Response Accuracy and B) Effect of Brand Attitude on Response Accuracy in case of Original Stimuli only.

### Reaction Times

The analysis of stimulus type on reaction times for both original and new stimuli revealed a significant impact of brand attitude on recognition memory of the original stimuli [t (38) = -4.55, p < .001, d = -0.73]. Figure 7 shows that the original stimuli were generally categorized in shorter time compared to the new ones (mean difference = -0.15 s).

**Figure 7.**
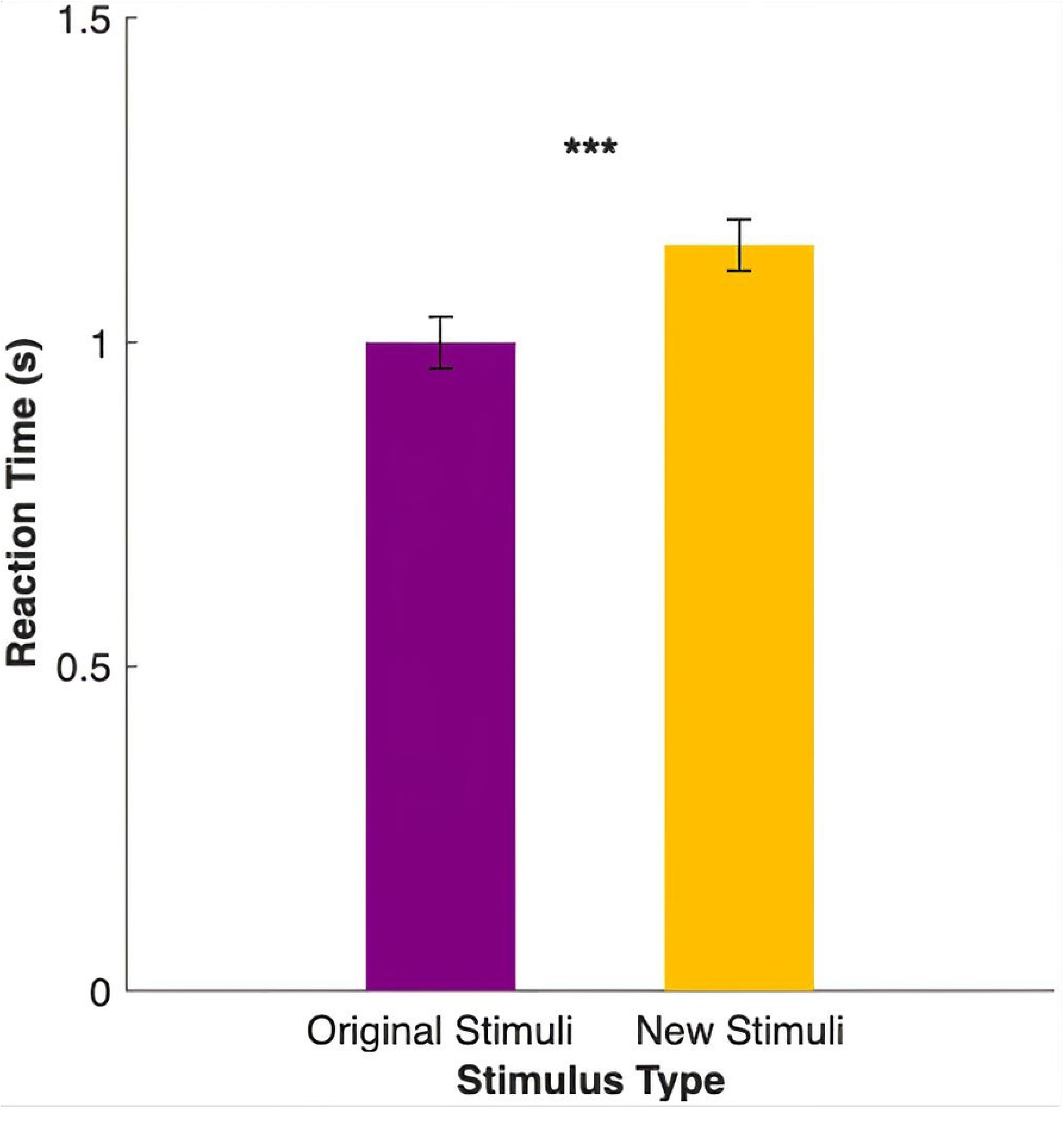
Effect of Stimulus Type on Reaction Times

However, no significant effect for brand attitude on reaction times in the original stimuli condition was observed [t (38) = -1.70, p = .097, d = -0.27]. Reaction times in both conditions were about 1 second, showing no effect of brand attitude on recognition speed.

## Summary

Taken together, brand attitude significantly influenced the accuracy of recognition memory of original stimuli, aligning with H3 suggesting that it has an effect on memory. However, brand attitude had no significant impact on reaction times.

## Discussion

This research aimed to establish a foundation for assessing how responses to brand activism are reflected in perceptual and cognitive mechanisms. We thus examined how consumer responses to brand activism manifest in brand attitudes, visual engagement, and memory performance. By integrating self-report, eye-tracking, and memory measures, the study advances the literature on brand activism, which by far has largely relied on surveys. The findings contribute in three key ways.

First, this study aligns with the asymmetry in consumer reactions observed by Mukherjee and Althuizen (2020), where disagreement with a brand’s sociopolitical stance produced significantly lower brand attitudes, while agreement did not deliver a corresponding positive effect. Results from our study indicate a distinct polarization of responses, with consumer-brand agreement associated with higher brand attitude and consumer-brand disagreement associated with lower brand attitude. This pattern highlights the heightened risks associated with brand activism and supports the notion that consumers increasingly expect brands to align with their values, making agreement a baseline expectation rather than a reward (18).

Second, the study contributes to research on visual engagement in consumer behavior. While overall time spent looking at advertisements was shaped by presentation order rather than agreement, eye-tracking revealed that disagreement elicited longer reading times for brand statements, reflecting deeper cognitive processing of dissonant information. Conversely, agreement increased fixations on brand-related elements (product and brand name), possibly consistent with approach–avoidance dynamics (19), for which stimuli with a positive valence generally elicit approach behaviors, whereas negative-valence stimuli are more likely to trigger avoidance responses. These findings extend theories of visual engagement (5, 7) to the domain of brand activism by showing how value alignment shapes perceptual engagement with marketing stimuli.

Third, the study demonstrates that brand attitudes influence memory performance. Positive brand attitudes enhanced recognition accuracy, indicating that agreement facilitates encoding through top-down mechanisms (16). This finding complements work on emotion and memory (10, 11, 20) by suggesting that positive brand alignment not only shapes brand evaluations, but also improves retention of advertising content. Linking these results with the eye movement behavior, a possible interpretation is that the more positive the participants’ attitude toward the brand, the more time they spend looking at it, and the better is their recognition of the product. This interpretation is further supported by the absence of an effect on reaction time, implying that the effect of familiarity on memory performance may occur during encoding or consolidation rather than through brand attitude.

Together, these contributions provide a more nuanced understanding of how brand activism affects consumer responses across conscious (attitudes) and less conscious (eye movements, memory) mechanisms, opening avenues for future research into the perceptual and cognitive mechanisms and the related neural underpinnings of consumer responses to activist messaging.

### 4.1 Managerial Implications

The results offer several prompts for marketers considering to include brand activism in their strategic plan. The evidence highlights an asymmetry: disagreement yields to strong negative effects, while agreement produces limited positive returns. Managers should recognize that activism may carry greater downside risks than upside potential, particularly when targeting heterogeneous audiences. When brands engage in activism, alignment with the values of their target audience is crucial. Misalignment can quickly erode trust and diminish brand equity, suggesting that activism is a viable strategy primarily when there is a clear fit with the brand’s identity and customer expectations. Moreover, the finding from this study that presenting activist messaging before advertisements reduces engagement with the ad underscores the importance of message sequencing. Brands may benefit from including activist messages within broader campaigns rather than placing them upfront, in order to preserve the main focus on the product. Ultimately, results on memory performance showed an association of positive brand attitudes with superior recognition of advertising content. This implies that when activism resonates with consumers, it can reinforce memory for brand messages and products, supporting long-term brand equity.

## Limitations

This study has several limitations that affect interpretation of the findings. First, the sample was unbalanced in terms of gender and distribution across agreement conditions, which may limit generalizability. Second, the use of AI-generated advertising images, while controlled for plausibility, may have reduced ecological validity, as some participants reported noticing artifacts (one participant reported that in some trials they were actively looking for AI-generated errors, while others noticed that certain details had an unrealistic appearance) and might have directed their attention to them. Third, participants’ self-reported attitudes were not validated with additional measures, raising the possibility of social desirability biases in categorization, also considering that a researcher was present in the lab throughout the experimental session. Finally, the laboratory setting may not fully capture naturalistic exposure to activism messages, where distractions and competing stimuli shape engagement.

## Future Research

Findings from this study pave the way for further investigations. First, future work should employ real brands and authentic campaigns to enhance ecological validity. Stronger personal involvement may amplify the observed effects on attitudes, attention, and memory. Second, research should more strongly consider naturalistic contexts, such as retail environments or social media feeds, where consumers encounter activism messages amid competing stimuli. An interesting perspective would be to also examine downstream behaviors such as purchase decisions or brand switching, providing direct evidence of how activism influences consumer choice and decision-making processes. Third, studies could manipulate visual features of advertisements (e.g., color, complexity) to disentangle the effects of activism from general design influences.

## Conclusion

Within consumer behavior research, it has been established that brand activism influences brand attitudes. This study demonstrates that brand activism influences consumer responses across multiple levels of processing. While alignment between consumer values and brand positions fosters positive attitudes, visual engagement, and memory, misalignment produces sharp declines in brand evaluation. The findings underscore both the promise and the risk of activism as a marketing strategy. For research, the study contributes new insights by integrating eye movement measures with attitudinal data. For managers, it highlights the need to carefully weigh risks, tailor causes to target audiences and consider how message sequencing affects consumer engagement. Ultimately, brand activism remains a high-stakes strategy: its success depends not only on what stance a brand takes, but also on how and to whom that stance is communicated.

## Acknowledgements

We would like to thank Dr. Bianca R. Baltaretu for her valuable feedback and consultation throughout the research process.

